# One more avenue for whale-watching contributions to science: the study of cetacean-cephalopod interactions

**DOI:** 10.1101/2021.04.23.440971

**Authors:** Stéphanie R.A. Suciu, Jasmine Zereba, Lorenzo Fiori, José M.N. Azevedo

**Affiliations:** Azorean Biodiversity Group (University of the Azores), Centre for Ecology, Evolution and Environmental Changes (CE3C), Rua da Mãe de Deus, 9500-321 Ponta Delgada, Portugal; Sea Colors Expeditions, 9500-764 Ponta Delgada, Portugal; Terra Azul Ltd., 9680-909 Vila Franca do Campo, Portugal

**Keywords:** cephalopods, DNA barcoding, teuthophagous cetaceans, whale watching, participatory science

## Abstract

Cephalopods are the primary source of food for several species of odontocetes. The unstable nature of this trophic resource is likely to affect the ecology of their cetacean predators, and this can be reflected on their conservation status but also on the tourist activities which target the observation of these animals. However, the study of the cetacean-cephalopod interaction is limited by the heavy logistics and expense of dedicated scientific campaigns. Fortunately, this gap can be filled by coupling modern molecular tools with indirect sampling methods. In this note we test if whale watching activities, which represent an intense observation effort worldwide, could be a source of material for studies of cetacean-cephalopod interactions. All contacted companies welcomed the invitation and received the sampling kit. Nine samples were collected, most of them in close association with sperm whales. All samples were determined as the seven-arm octopus *Haliphron atlanticus* (Octopoda, Alloposidae). We conclude that, although the Azores may have particularly favorable conditions for participatory science, similar programs can be replicated elsewhere

---

Cephalopods have a structuring role in oceanic ecosystems, acting as a link between different trophic levels by feeding on a variety of fish and invertebrates and being in turn eaten by a wide range of predators including fish, sharks, marine mammals and seabirds (de la Chenais et al., 2019; Escánez et al., 2020). The short life span of most species accounts for variable population sizes over time, their population dynamics tracking environmental conditions, such as food abundance or water temperature (Forsythe, 2004; Jackson and Moltschaniwskyj, 2002). The resulting variability in cephalopod biomass can directly affect the biomass of their prey and can also influence their predators, including marine mammals (Liu & Chen, 2009). The unstable population dynamics of oceanic cephalopods is likely to be a key factor influencing the distribution of teuthophagous cetaceans, with effects on conservation but also economic given the importance of whale watching (O’Connor et al., 2019).

Because cephalopod habitats, behaviors, morphology and life-history strategies vary so greatly, a single standard approach does not suffice. Direct sampling methods include trawling and video-surveys (Hoving et al., 2014). However, each of these has its own biases. The sophisticated features associated with the predatory behavior of cephalopods such as vision and agility, for example, may result in avoidance behavior towards many types of oceanographic gears (Villanueva et al., 2017). On the other hand, even specimens captured in trawled nets may be damaged, making their morphological identification challenging (Vecchione et al., 2010).

An indirect method of studying cephalopods and determining their presence in a region, is by analysis of remains found in the stomach content of their predators (Clarke et al., 1993). Being crucial to look at cetacean diet, this method has limitations for studying cephalopod ecology, stemming from predator selectivity and unprecise location data. Even so, it has been claimed that stomach content analysis provides a better overview of cephalopod distribution and relative abundance than net catches (Clarke, 2006). However (and fortunately) research on cetacean diet from stomach contents almost stopped with the 1980’s IWC moratorium on whale hunting. Some of the methodological, logistic, and ethical limitations to sampling pelagic cephalopods can nevertheless be overcome with molecular approaches (O’Brien et al., 2018). DNA barcoding, for instance, allows individual species identification based on an organism’s remains but also to determine community composition from DNA fragments persisting in environmental samples, such as water (Valentini et al., 2009). Using these methods, species can be determined from damaged samples captured by commercial nets (e.g. Dai et al., 2012), and partially digested remains from seabirds, fishes, stranded whales and seals stomachs (Hoving et al., 2014; Xavier et al., 2015).

Cephalopod remains are occasionally observed at the surface in connection with cetacean feeding activities (Fig. 1), and they have been used inconsistently as a source of information on deep water cephalopods (Clarke, 1996c). We could not find publications using this material, which is not surprising because of its unpredictable source which, to be systematically collected, would require an intense and targeted effort, not compatible with the standards of regular scientific work. Nevertheless, we did find references to the observation of cephalopod remains in the context of whale watching activities (Escanez et al. 2019; Sarabia-Hierro & Rodriguéz-González, 2019). Whale watching is an important tourist industry worldwide, and it represents a huge observation effort in terms of number of boats, miles crossed, and hours spent at sea (O’Connor et al., 2009), unmatched by standard scientific campaigns. Harnessing this effort can provide inexpensive and valuable information about marine life, particularly in data deficient areas. Examples of the scientific use of opportunistic data from whale watching include the study of coastal cetaceans off the southwest coast of South Africa (Vinding et al. 2015) or the analysis of how temporal scales influence cetacean ecological niche modelling (Fernandez et al. 2018). We therefore argue that whale watching operations worldwide, with their intense observation effort often targeted at teuthophagous cetaceans, are ideal platforms to collect floating cephalopod remains and make them available for scientific purposes. The present paper presents the results of a pilot project set up to test this idea.

**Fig. 1.**
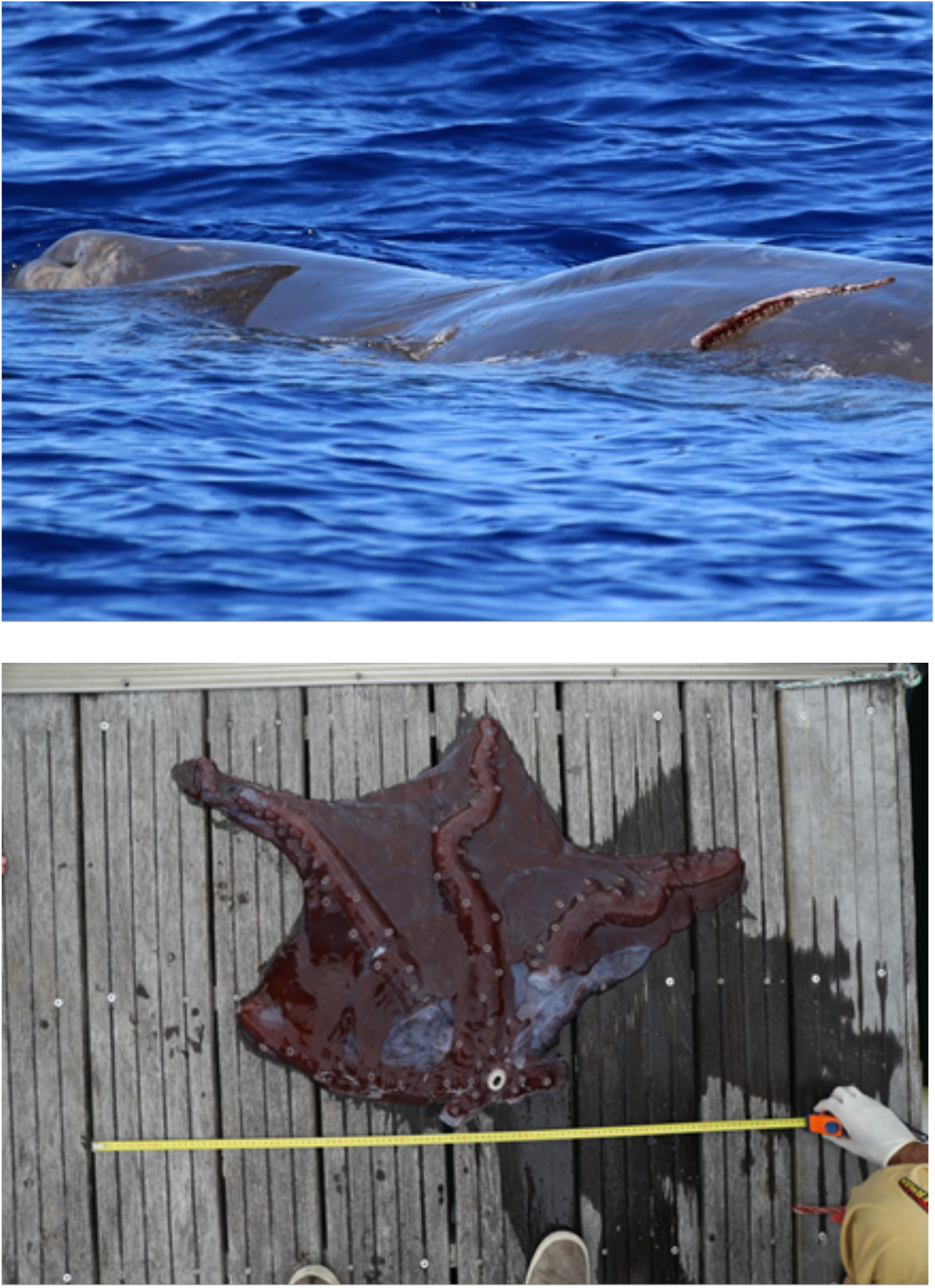
Examples of remains of cephalopods found during whale watching activities. A tentacle on a sperm whale (fig.1(a), credit: Sea Colors Expeditions) and *Haliphron atlanticus* remains photographed on shore (fig.1(b), credit: Terra Azul)

The study was carried out in 2020, on São Miguel Island, Azores, where a whale watching industry is operating since the early 1990’s (Silva, 2015). All the five companies operating in that year accepted to participate. Each company received a boxed sampling kit, and staff training on the collection of samples and related information. The protocol was simple: every time a vessel would spot cephalopod remains they would collect a piece (including the beak, if present) and preserve it in 96% ethanol inside the supplied jars. Additionally, they were requested to collect, as a minimum, the time and geographic position, and information about the species of cetaceans in the area and their behavior. If possible, photographs of the remains should also be taken. Upon return to base a contact would be made with the research team which would pick up the samples and store them at 4°C. The project was cost-free for the companies, and was run on a 300€ budget (not including the human resources cost). A transparency and open data policy was implemented from the beginning, with a website (https://moniceph.wixsite.com/home) set up so that the companies and the public could follow the project activities and results.

At the end of the tourist season the samples were sent to an external laboratory which took care of DNA extraction and PCR amplification of the mitochondrial COI gene using the primers jgHCO2198 and jgLCO1490 (Geller et al., 2013). Forward and reverse sequences were assembled using Geneious (v. R10, Biomatters, Auckland, NZ) and reciprocally verified to generate a complete contig of the sequenced fragments. All contigs were compared to the BOLD reference database and the NCBI nucleotide database using the BLAST algorithm (https://blast.ncbi.nlm.nih.gov/Blast.cgi) for taxonomic assignment. The images collected were also analyzed for morphological species determination.

A total of 9 samples were collected, including some taken the previous year. Information about an additional finding for which a biological sample was not available was also recorded. This information, and the results of the morphological and molecular species determination, are given in Table 1. The seven-arm octopus *Haliphron atlanticus* (Octopoda, Alloposidae) was the only species identified. On most occasions, sperm whales were present in the area where remains were found. Although Clarke et al. (1993) reported that sperm whales in the Azores feed primarily on Octopoteuthidae, Histioteuthidae and Architeuthidae squids, *H. atlanticus* is also part of this specie’s diet (see, recently, Cherel et al., 2021). Common dolphins were associated with the two samples without the presence of sperm whales.

**Table 1.**
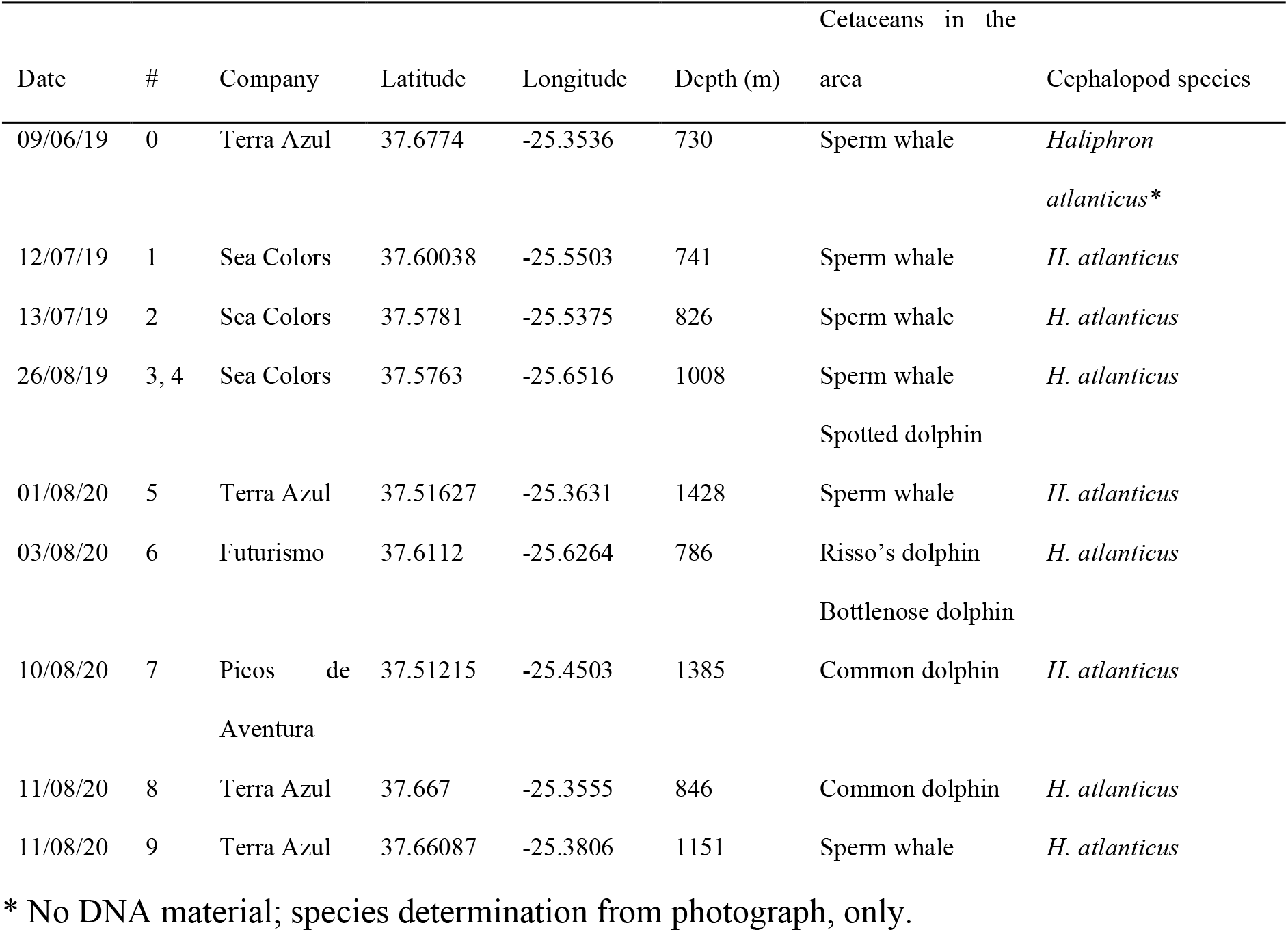
Cephalopod samples collected, associated information and species determination by DNA barcoding and image analysis, except where otherwise stated.

The opportunistic nature of samples collected in this manner is likely to introduce bias. The fact that only samples of *H. atlanticus* were collected could, for instance, be due to the “watery” nature of this species, with the predator focusing mainly on the buccal mass and rejecting the rest, as speculated by Santos et al. (2001) for blue sharks. Carried out consistently across time and space a monitoring program such as the one presented here could clarify these biases while providing relevant information on issues like changes in relative abundance of oceanic cephalopods or the diet of their cetacean predators.

In summary, our results show that, by collecting remains of cephalopods associated with cetacean observations, whale watching companies can contribute to fill a gap in indirect research on cephalopod ecology and cephalopod-cetacean interactions.

In our experience, companies showed immediate interest for the project and have expressed their willingness to continue collaborating with it. We believe this attitude is based on the low cost of this collaboration compared to its perceived added value, not only to the marketing image of the company (resulting from being associated with research activities) but also to the improvement of client experience. This is in line with the results of Bentz et al. (2016) which demonstrate that whale watchers in the Azores value receiving information about the cetaceans and their environment. But another important factor was the motivation demonstrated by the tour guides which, for the most part, had degrees on biological sciences because this is a legal requirement. Thus, it may be that the institutional conditions in the Azores are favorable to the engagement of the industry with participatory science (e.g. Philips et al., 2019; Wuebben et al., 2020). In retrospect, however, we seem to have followed most of the engagement framework laid out by Pandya (2012): our request was aligned with community priorities, we built on existing knowledge and validated previous practices and we made a point of disseminating the results widely, so that effort was recognized. We therefore believe that our results are replicable elsewhere, and encourage the establishment of communities of practice between members of the whale watching and the research communities, in order to take full advantage of this research opportunity.

## Acknowledgements

We appreciate the positive and encouraging response obtained from the managers and staff of the whale watching companies participating on this pilot project: Sea Colors Expeditions, Terra Azul, Futurismo, Moby Dick, Picos de Aventura. We thank Jasmine Zereba, Lorenzo Fiori, Marylou Féat, Mariana Silva, Ana Castanheira and Vanessa Costa for collecting the samples while operating whale watching activities.

